# Age-Stratified Meta-Transcriptomic Analysis Reveals Early and Core Dysregulated Pathways in Parkinson’s Disease

**DOI:** 10.64898/2026.07.14.738591

**Authors:** Prakashini Saroj Nilgirwar, Sahil Jain, Bhakti Badhe, Rituja Shinde, Manoj Baranwal, Dimple Davray

**Affiliations:** Bioinformatics Research Centre, Dr. D.Y. Patil Biotechnology and Bioinformatics Institute, Dr. D.Y. Patil Vidyapeeth, Pune, Maharashtra, India; Bioinformatics Centre, Savitribai Phule Pune University, Pune, Maharashtra, India; Center of Excellence in Emerging Diseases, Department of Biotechnology, Jaypee Institute of Information Technology, Noida, Uttar Pradesh, India; Department of Biotechnology, Thapar Institute of Engineering and Technology, Patiala, 147004 Punjab, India; Medical Research Council Unit, The Gambia at London School of Hygiene and Tropical Medicine, The Gambia

**Keywords:** Parkinson’s disease, Age-stratified transcriptomics, Bulk RNA-seq meta-analysis, Early-onset Parkinson’s disease, Biomarker discovery, Neurodegeneration

## Abstract

Parkinson’s disease (PD) is strongly age-associated, yet how aging reshapes PD-related transcriptional changes remains unclear. We performed an age-stratified meta-analysis of 16 bulk RNA-seq datasets (646 samples: 314 PD, 332 controls) to distinguish early-onset (<60 years) from late-onset (≥60 years) signatures. In the full dataset, 131 significantly (padj<0.05, |log2FC|>1) differentially expressed genes (DEGs) were observed, spanning neuronal activity-dependent genes (NPAS4, PVALB, ARC, FOSB) and immune/stress-related transcripts (IL3RA, SLC25A6, HSPA1A/B). Age-specific analyses revealed 31 DEGs in <60, dominated by large-effect changes in uncharacterized lncRNA LINC02188, pseudogene loci (MUC20P1, RPS28P7), calcium-modulating gene CALML6, lipid□associated gene TLCD3B, and cytoskeletal regulators (TIAM2, KCTD8). In contrast, the ≥60 group showed 181 DEGs enriched for neuronal markers (NPAS4, PVALB) and immune–metabolic genes (FGA, NPC1L1, UPK1A, HSPA1A/B). GO/KEGG analyses indicated that the <60 signature centers on actin remodeling, filopodia, axonogenesis, and Rap1-mediated adhesion/signaling, consistent with early neurite and structural reorganization. The ≥60 signature was enriched for blood microparticles, chemokine activity, infection-related pathways, ER protein processing, and arachidonic/ether lipid and cytokine signaling, pointing to broad immune–metabolic and proteostasis dysregulation. Cross-age comparison showed that classical PD neuronal immediate-early gene changes are largely ≥60-driven, whereas early-onset PD involves novel lncRNA-calcium-lipid and cytoskeletal modules. These findings highlight LINC02188, TLCD3B and related cytoskeletal/lncRNA genes as novel early-onset PD-associated candidates, and NPC1L1, IL3RA and PVALB as age-amplified markers within the broader PD transcriptomic signature.

## 1. Introduction

Parkinson’s disease (PD) is the second most prevalent neurodegenerative disorder, following Alzheimer’s disease, and is characterized by chronic inflammation, aggregation of α-synuclein, immune dysfunction, and loss of dopaminergic neurons in the substantia nigra pars compacta (Bloem et al., 2021). This leads to non-motor symptoms, such as cognitive impairment, psychiatric disturbances (including depression and anxiety), and dysfunction in the autonomic nervous system, alongside motor symptoms, such as rest tremors, bradykinesia, rigidity, and postural instability (Silva et al., 2023). Among PD individuals, around 50 percent are affected by anxiety and/or depression at some stage of illness, and this is linked to elevated risk of dementia, cognitive and physical decline, and higher mortality rates (Burchill et al., 2024).

Aging populations contribute significantly to the rise in PD cases, as characterized by the increase in crude prevalence and age-standardized rates by 74% and 22% respectively during 1990-2016 (Dorsey et al., 2018). Indeed, PD affects approximately 0.3% of the world population (∼11.7 million registered PD cases in 2021), but the percentage rises to 4% in those >80 years of age (Luo et al., 2025). PD cases are projected to reach 25.2 million by 2025, primarily driven by an aging population, with age-standardized prevalence expected to increase by 55% and all-age prevalence projected to increase by 76% between 2021 and 2050 (Su et al., 2025). Despite this strong age-dependence, how aging reshapes the molecular landscape of PD remains unclear.

PD is diagnosed clinically as per the Movement Disorder Society (MDS) criteria, which is based on the presence of symptoms such as tremors, bradykinesia, and rigidity (Berg et al., 2018). The clinical diagnosis of PD is around 80% accurate, and is only confirmed by postmortem brain examination (Yamashita et al., 2023). Early symptoms often overlap with other neurodegenerative or psychiatric disorders, thus, resulting in frequent misdiagnosis. Therefore, the identification of reliable biomarkers is important as it may improve diagnostic precision as well as enable early detection, consequently facilitating development of early disease-modifying therapies (Chen-Plotkin et al., 2018).

Recent studies have tried to identify reliable biomarkers for early Parkinson’s disease (PD). Xiang et al. (2022) identified cerebrospinal fluid (CSF) markers, such as total α-synuclein and neurofilament light chain, that help differentiate PD from atypical parkinsonian syndromes (Xiang et al., 2023). A 2024 study reported that circulating microRNAs (such as miR-30c, miR-331-5p), particularly miRNA clusters, exhibit a high diagnostic accuracy despite variations in detection protocols (Zhang, W. et al., 2024). More recently, 46 blood and urine biomarkers were identified, linking the onset of PD to systemic inflammation and metabolic dysregulation, with sex-specific and age-specific differences (Gao et al., 2025). Together, these studies present promising biomarker candidates, and also highlight the complexity and heterogeneity of PD pathophysiology.

However, these approaches are limited by methodological heterogeneity (Xiang et al., 2023), population bias, and lack of longitudinal validation in pre-symptomatic stages (Zhang, W. et al., 2024). This may lead to an overestimation of diagnostic accuracy. Moreover, majority recent studies do not explicitly account for age-dependent molecular variation, although aging is the strongest risk factor for PD. To address these gaps, we performed a meta-analysis of RNA-seq datasets from PD patients and healthy controls, and stratified them into early-onset (<60 years) and late-onset (≥60 years) groups. We applied differential expression, functional enrichment, and network-based analysis, in order to identify age-specific and core transcriptional signatures that reflect the molecular mechanisms of PD progression and may serve as early biomarkers or therapeutic targets.

## 2. Material and Methods

### 2.1. Data Collection

The NCBI Gene Expression Omnibus (GEO) database was used to download the transcriptomic data with the search keywords “Parkinson”s” and “Homo sapiens”. Only studies categorized as either “Expression profiling by array” or “Expression profiling by high throughput sequencing” were selected to ensure consistency with transcriptomic profiling methodologies. Duplicate entries were removed, and the remaining 79 transcriptomic studies spanning across five continents were included for downstream analysis.

### 2.2. Data Preprocessing and Exploratory analysis

Metadata files corresponding to the shortlisted studies were downloaded and pre-processed by organizing the data structure, addressing missing values, and standardizing variable formats. Exploratory data analysis was performed using R packages including dplyr, tidyverse, and ggplot2. The samples were categorized based on their study type and geographical distribution. Further, age distribution across different countries was plotted and statistically assessed by one-way analysis of variance (ANOVA), followed by Tukey”s Honest Significant Difference (HSD) post hoc test for pairwise comparisons.

### 2.3. Differential Gene Expression Analysis

Based on the availability of raw FASTQ files, 16 bulk RNA-seq studies (from North America, Asia, and Europe) were selected for Differential Gene Expression (DGE) analysis. Studies either lacking raw data or involving altered glucose levels, brain stimulation condition and/or samples from stem cells were excluded. Quantification was carried out using the OneStopRNAseq online platform (Li et al., 2020), resulting in a count matrix for each study. Further analysis was performed in R, retaining genes expressed in at least three samples. The entire DGE workflow, including batch correction, normalization, and differential expression analysis, was performed using the dplyr, sva, limma, and DESeq2 packages. Initially, DGE analysis was conducted across all samples to identify globally differentially expressed genes (DEGs). Later, samples were grouped by age (<60 and ≥60 years), and group-wise DGE analysis was performed to identify age-associated gene expression changes and potential biomarkers. To prioritize biologically meaningful differentially expressed genes (*in lieu* with current biomarker discovery frameworks), we selected top candidates based on absolute log2 fold change (|log2FC|>1) following filtering by adjusted p-value (padj < 0.05).

### 2.4. Data Visualization

Global study distribution was visualized by mapping the number of transcriptomic studies per country. Metadata were aggregated in R, and country boundary shapefiles from rnaturalearth were imported as sf objects. Study counts were joined to the spatial features using standard geospatial merges. Data processing and visualization were performed with dplyr and ggplot2, with countries shaded on a continuous gradient by study count.

Gender-based age analysis was performed in R using dplyr, tidyverse, and ggplot2. Samples with available age and gender annotations were plotted as box-and-whisker distributions with overlaid data points. For geographic age analysis, samples with age and country information were grouped by country, and age distributions were visualized using country-wise boxplots. Differences in mean age across countries were tested using one-way ANOVA, followed by Tukey”s HSD post hoc test for pairwise comparisons.

Volcano plots were generated using GraphPad Prism to identify and visualize the significant differentially expressed genes (DEGs) across all samples, as well as across the age-groups (<60 and ≥60 years). The top candidates (padj < 0.05; |log2FC|>1) were color coded as blue (downregulation) and red (upregulation).

### 2.5. Functional Enrichment analysis

Gene Ontology (GO) and Kyoto Encyclopedia of Genes and Genomes (KEGG) pathway analysis were conducted with clusterProfiler and org.Hs.eg.db packages in R, to determine the highly enriched biological processes, molecular functions, cellular components, and signaling pathways. All analysis were conducted with an adjusted p-value threshold of < 0.05 (Benjamini-Hochberg correction) to identify statistical significance.

## 3. Results

### 3.1. Dataset Collection and Preprocessing

As of June 2024, the NCBI Gene Expression Omnibus (GEO) contained 4,316 studies related to gene expression in human Parkinson”s disease (PD) (Supplementary Sheet S1). 130 studies classified under the “Expression profiling by array” and 170 studies classified under the “Expression profiling by high throughput sequencing” were selected as they are standardized platforms providing gene expression data for integrative analysis. Amongst these 300 studies, reports based on genomic data, *in vitro* cell line experiments, and other neurodegenerative disorders were filtered out. Lastly, duplicate studies were removed, and a total of 79 relevant studies were selected (Supplementary Sheet S2). These included 22 bulk mRNA studies, 4 miRNA sequencing studies, 1 non-coding RNA study, 5 single-cell RNA-seq studies, and 47 microarray-based studies (Table 1). The shortlisted dataset consisted of diverse tissue sources, including 1,739 blood samples, 1,681 brain-derived samples, 77 skeletal muscle samples, 64 skin samples, 59 appendix samples, 46 stem cell-derived samples, and 30 colon samples. A total of 3,000 control samples and 1,745 PD samples were collected from five continents: North America (33 studies), Europe (29), Asia (14), Oceania/Australia (1), and South America (2), as detailed in Table 1. The country-level distribution of studies and samples is shown in Fig. 1 and Fig. 2 respectively.

**Fig. 1.**
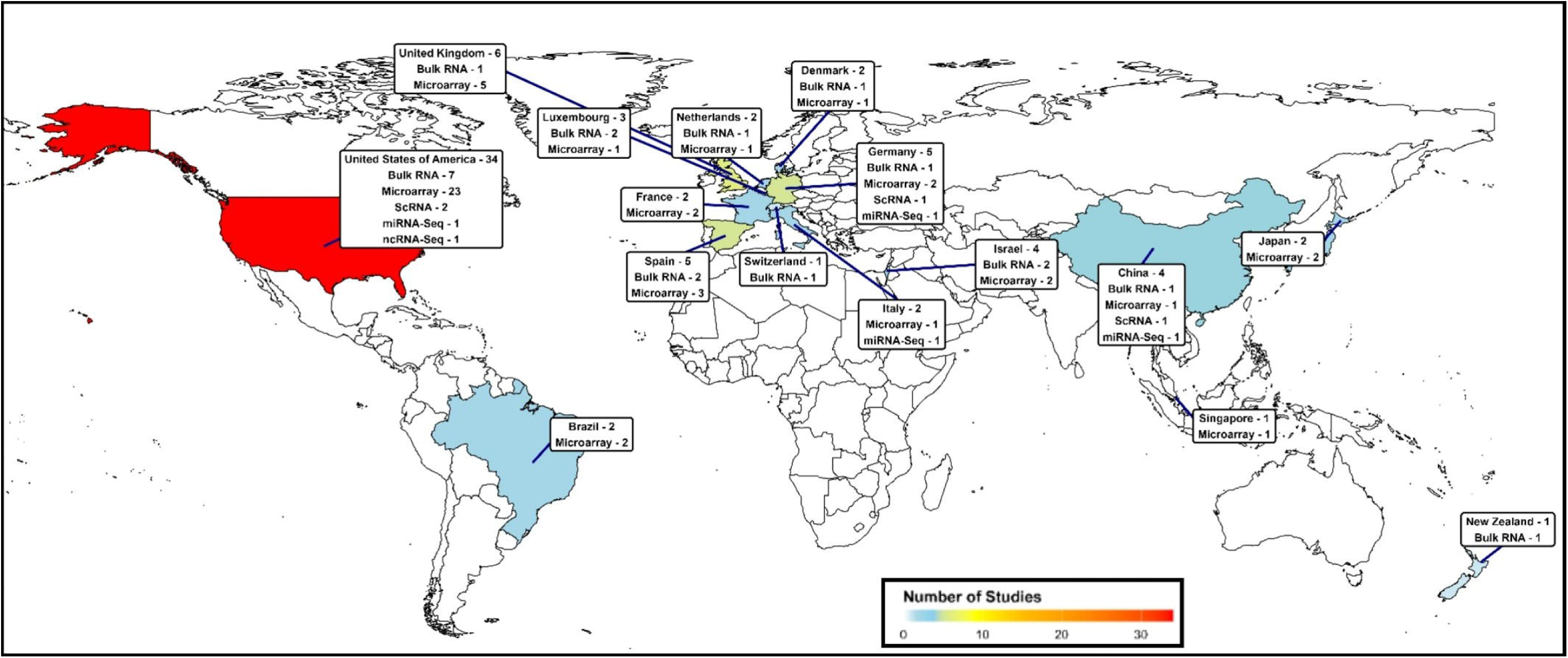
Global distribution of the transcriptomic studies. The color gradient (blue to red) represents the number of studies per country, with red indicating the highest count. Labels show the total studies per country and the number of studies by sequencing types, including bulk RNA, Microarray, ScRNA, miRNA-seq, and ncRNA-seq studies.

**Fig. 2.**
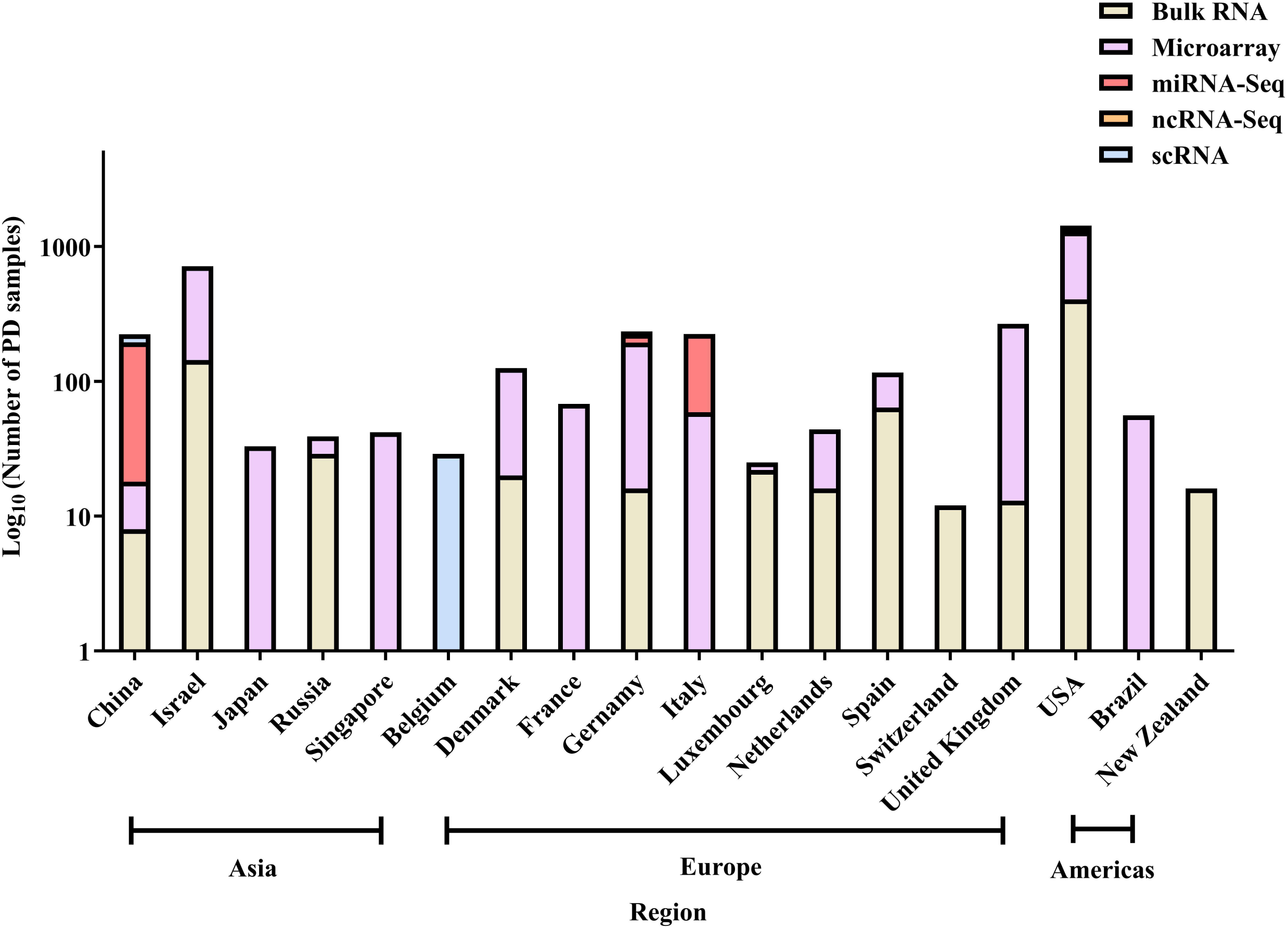
Distribution of Parkinson”s disease samples across countries, categorized by RNA sequencing platform. Bars represent the number of samples obtained using five distinct sequencing technologies: Bulk RNA sequencing (red), Microarray (green), miRNA-Seq (teal), ncRNA-Seq (blue), and ScRNA-Seq (purple). The y-axis is shown on a log_10_ scale to account for the large differences in sample counts between countries and to enable clearer visual comparison across regions.

**Table 1.** Overview of the transcriptomic studies included in the exploratory analysis, categorized by study type, geographic distribution, and total control and PD samples.

| Study Type | Number of studies in each continent |  |  |  |  | Total number of studies | Total Control samples | Total Disease samples |
| --- | --- | --- | --- | --- | --- | --- | --- | --- |
|  | North America | Europe | Asia | Australia | South America |  |  |  |
| **Bulk RNA** | 7 | 9 | 5 | 1 | - | 22 | 386 | 379 |
| **Microarray** | 22 | 16 | 7 | - | 2 | 47 | 2337 | 1074 |
| **miRNA** | 1 | 2 | 1 | - | - | 4 | 215 | 212 |
| **Non-coding RNA** | 1 | - | - | - | - | 1 | 14 | 14 |
| **Single cell RNA** | 2 | 2 | 1 | - | - | 5 | 48 | 66 |
| **Total** | **33** | **29** | **14** | **1** | **2** | **79** | **3000** | **1745** |

### 3.2. Exploratory and Statistical Analysis

Amongst the 79 shortlisted studies, the bulk mRNA studies (22) were considered for further analysis to ensure consistency across datasets. However, 6 bulk mRNA studies involved samples collected under conditions such as altered glucose levels, brain stimulation, or from PD carriers, and were therefore excluded from further analysis. Overall, 16 bulk mRNA studies, distributed amongst 3 continents and 8 countries, were finally considered (Supplementary Sheet S3; Table 2). A total of 332 control and 314 PD samples were found in these studies (Table 2).

**Table 2.**
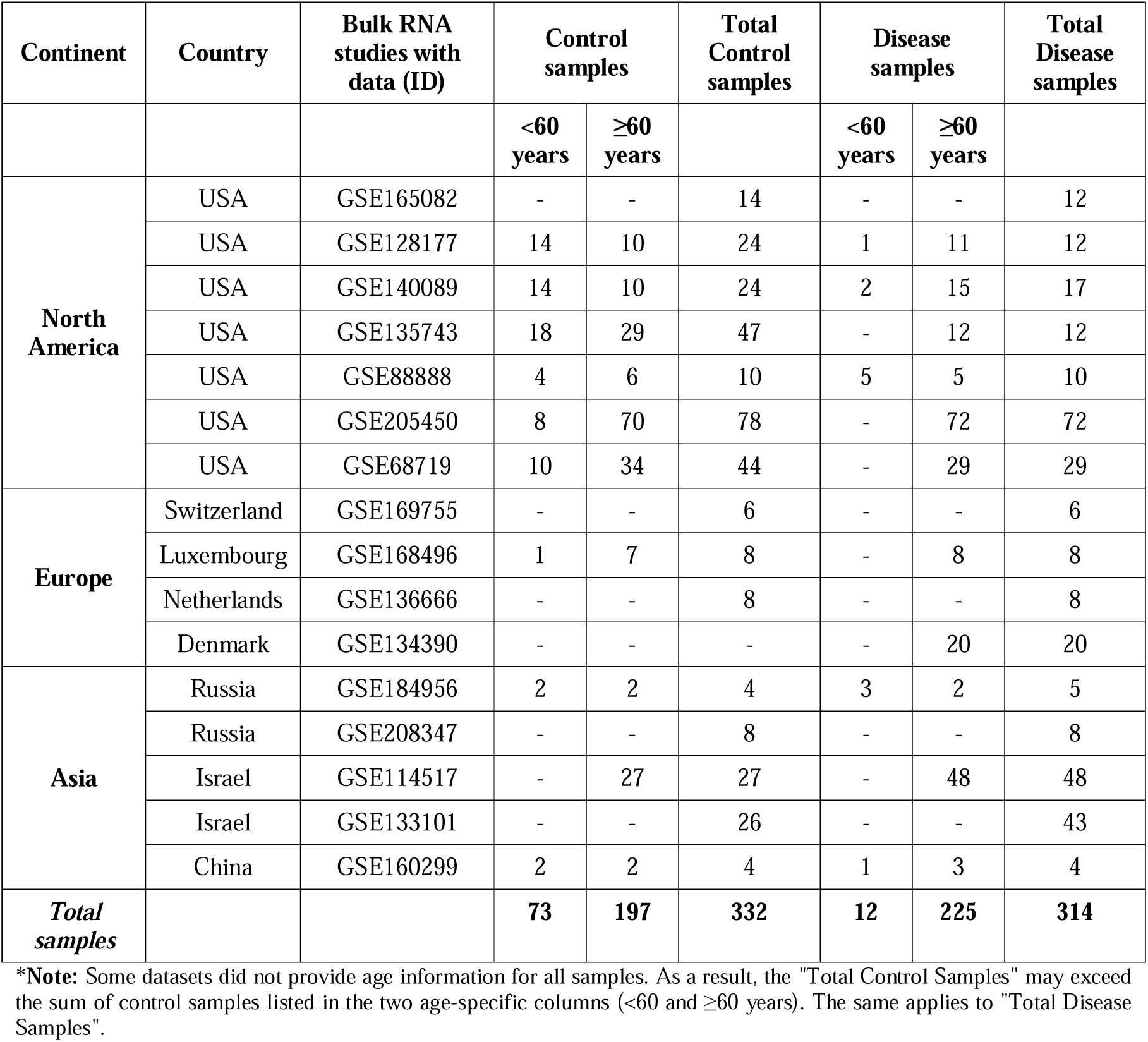
Summary of bulk RNA-seq datasets across countries by age group and disease status. The table reports the number of control and PD samples stratified into two age groups: <60 years and ≥60 years, along with the total number of control and disease samples per study.

| Continent | Country | Bulk RNA studies with data (ID) | Control samples |  | Total Control samples | Disease samples |  | Total Disease samples |
| --- | --- | --- | --- | --- | --- | --- | --- | --- |
|  |  |  | <60 years | ≥60 years |  | <60 years | ≥60 years |  |
| North America | USA | GSE165082 | - | - | 14 | - | - | 12 |
|  | USA | GSE128177 | 14 | 10 | 24 | 1 | 11 | 12 |
|  | USA | GSE140089 | 14 | 10 | 24 | 2 | 15 | 17 |
|  | USA | GSE135743 | 18 | 29 | 47 | - | 12 | 12 |
|  | USA | GSE88888 | 4 | 6 | 10 | 5 | 5 | 10 |
|  | USA | GSE205450 | 8 | 70 | 78 | - | 72 | 72 |
|  | USA | GSE68719 | 10 | 34 | 44 | - | 29 | 29 |
| Europe | Switzerland | GSE169755 | - | - | 6 | - | - | 6 |
|  | Luxembourg | GSE168496 | 1 | 7 | 8 | - | 8 | 8 |
|  | Netherlands | GSE136666 | - | - | 8 | - | - | 8 |
|  | Denmark | GSE134390 | - | - | - | - | 20 | 20 |
| Asia | Russia | GSE184956 | 2 | 2 | 4 | 3 | 2 | 5 |
|  | Russia | GSE208347 | - | - | 8 | - | - | 8 |
|  | Israel | GSE114517 | - | 27 | 27 | - | 48 | 48 |
|  | Israel | GSE133101 | - | - | 26 | - | - | 43 |
|  | China | GSE160299 | 2 | 2 | 4 | 1 | 3 | 4 |
| **Total samples** |  |  | **73** | **197** | **332** | **12** | **225** | **314** |

**\*Note:** Some datasets did not provide age information for all samples. As a result, the "Total Control Samples" may exceed the sum of control samples listed in the two age-specific columns (<60 and ≥60 years). The same applies to "Total Disease Samples".

To analyze age-specific patterns, the samples were segregated into two classes based on age groups viz., <60 and ≥60 years (Table 2). Samples with undefined age were included in overall analysis but excluded from age-stratified comparisons. Statistical analysis revealed no significant difference in age between female and male participants (p = 0.115), indicating that age was relatively similar across genders within the PD samples (Fig. 3 a)). In contrast, a highly significant difference in age distribution was observed among participants from different countries (p = 2.05 × 10□□; Fig. 3 b)), indicating a significant relevance of conducting age-specific analysis across geographic locations and possibility of region-specific early age PD biomarkers.

**Fig. 3.**
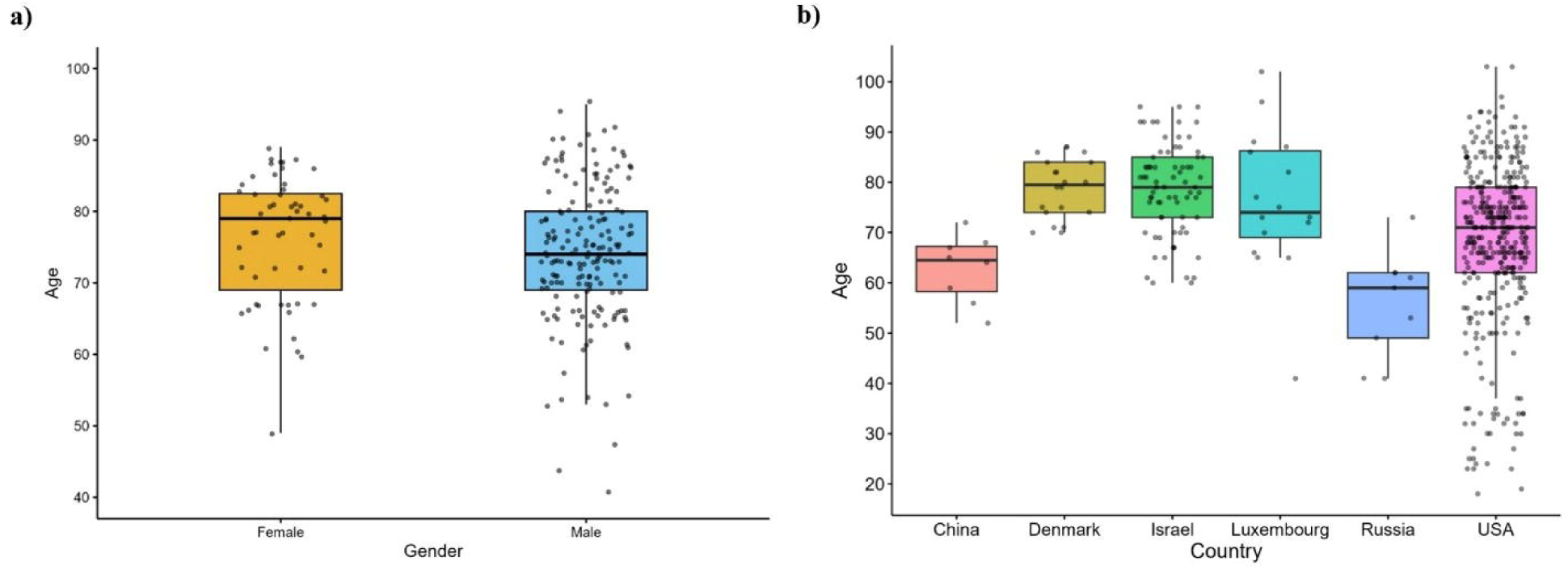
Boxplots illustrating a) The age distribution of Parkinson”s disease samples stratified by gender (n = 55 for Female, n = 178 for Male). Female and male participants are color-coded in orange and blue, respectively. b) Age distribution of samples across six countries: China (n = 8), Denmark (n = 20), Israel (n = 75), Luxembourg (n = 16), Russia (n = 9), and USA (n = 379). Countries with no available age data (Netherlands and Switzerland) were excluded. Each box represents the interquartile range (IQR), with the horizontal line indicating the median.

### 3.3. Bulk RNA-Based Differential Gene Expression Analysis

An initial DGE analysis was performed on the complete dataset to identify genes commonly dysregulated in PD across all ages (Supplementary Sheet S4). To investigate age-specific gene expression and potential early biomarkers, the complete dataset was segregated based on age-group (<60 and ≥60 years) and separate DGE analysis were then conducted (Supplementary Sheet S5, S6). It is to be noted that a transcript was considered to be significantly dysregulated only if the padj-value < 0.05 and a |log2 fold change| > 1 was observed.

#### 3.3.1. Differential gene expression across the complete dataset

DGE analysis of the complete dataset (n=646; 332 controls, 314 PD) identified 131 significantly dysregulated genes, *assumedly* representing a biologically meaningful transcriptional signature of PD across all ages. The top 20 downregulated genes were led by SLC35D3 (a Golgi nucleotide□sugar transporter; log2FC −1.476), MMP3 (a matrix metalloproteinase; −1.469), CCL3L3 (an inflammatory chemokine; −1.323), CHRNA2 (a nicotinic acetylcholine receptor subunit; −1.278), and neuronal activity genes including NPAS4 (-1.275), FOSB (-1.261), PVALB (-1.256), and ARC (-1.220), alongside EGR family members (Fig. 4 a)). This combination of impaired vesicle/matrix remodeling, chemokine signaling, cholinergic transmission, and immediate-early gene expression points to a coordinated suppression of activity□dependent neuronal and neuroimmune programs (Fan and Xiao, 2018; Shepard et al., 2019).

**Fig. 4.**
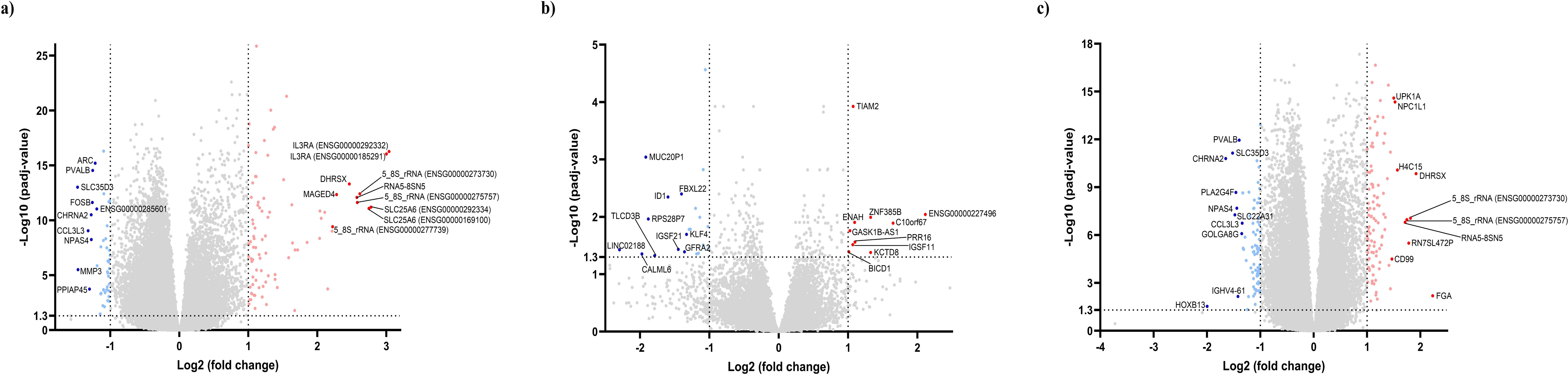
Volcano plots of differentially expressed genes (DEGs) in Parkinson”s disease. a) Plot from all the samples from 16 studies, b) Plot for age group <60 years across the 16 studies, and c) Plot for age group ≥60 years across the 16 studies. Each dot represents a gene, plotted by log2 fold change (x-axis) and –log_10_ adjusted p-value (y-axis). Genes with adjusted p-value < 0.05 and |log2 fold change| > 1 were considered significantly differentially expressed. The downregulated, upregulated, and non-significant genes are color coded blue, red, and grey respectively. The top 10 dysregulated genes are labelled in all the graphs. When all the datasets were analyzed together, 131 genes were significantly dysregulated (44 downregulated, 87 upregulated). Stratification by age revealed 31 dysregulated genes in individuals <60 years (21 downregulated, 10 upregulated) and 181 dysregulated genes in individuals ≥60 years (80 downregulated, 101 upregulated), underscoring that the overall PD signature is largely driven by the ≥60 group and that age strongly shapes the detectable transcriptional response.

In contrast, the top 20 upregulated genes were dominated by immune-associated transcripts like IL3RA (3.039), SLC25A6 (2.776), and non-coding RNAs including multiple 5_8S_rRNA species (2.615) (Fig. 4 a)), reflecting mitochondrial stress and immune activation. This pattern suggests heightened peripheral immune responses alongside neuronal suppression, consistent with systemic PD biology (Kung et al., 2022; Roodveldt et al., 2024).

#### 3.3.2. Differential gene expression based on age

The age-stratified dataset comprised 237 PD samples and 270 control samples, distributed as follows: <60 group – 73 control and 12 PD samples; ≥60 group – 197 control and 225 PD samples (Table 2). Amongst the 73 control samples in the <60 group, 37 control samples were excluded as they corresponded to studies wherein no PD sample was defined for age <60. Therefore, the total number of control and PD samples considered for the DGE analysis of the <60 group were 36 and 12 respectively.

*For the <60 age-group* - DGE testing of 42,816 filtered genes identified 31 significantly dysregulated genes, comprising 21 downregulated and 10 upregulated genes (Fig. 4 b)). The top downregulated genes were novel candidates LINC02188 (log2FC = -2.296), CALML6 (-1.973), MUC20P1 (-1.916), RPS28P7 (-1.880), and TLCD3B (-1.789), with no prior PD associations (Table 3). Upregulated genes included KCTD8 (1.325), PRR16 (1.100), and TIAM2 (1.073), suggesting early neuronal remodeling (Schwenk et al., 2016; Tolias et al., 2011) (Table 3). This focused signature of largely novel genes suggests distinct early-onset PD biology.

**Table 3.**
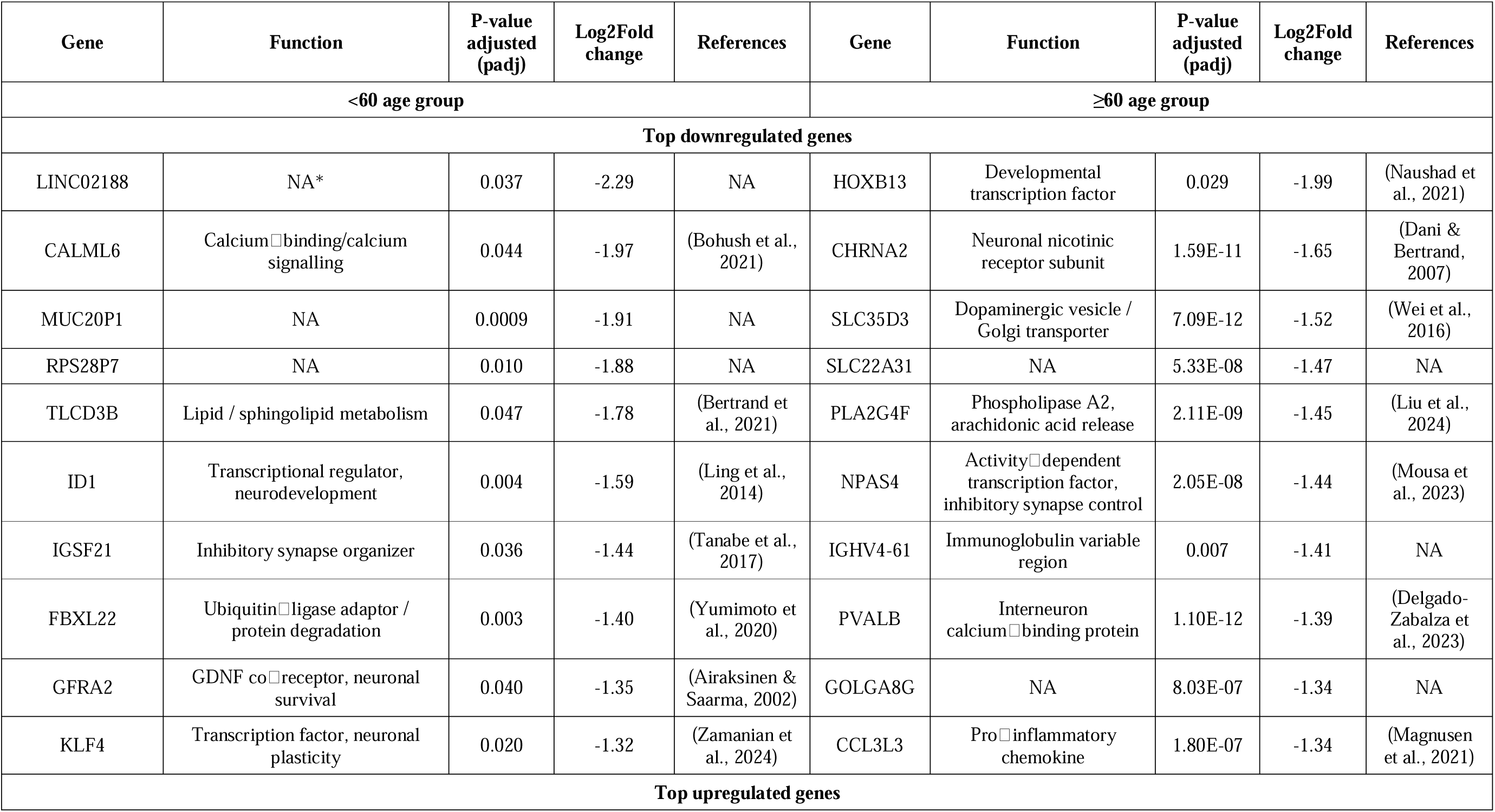
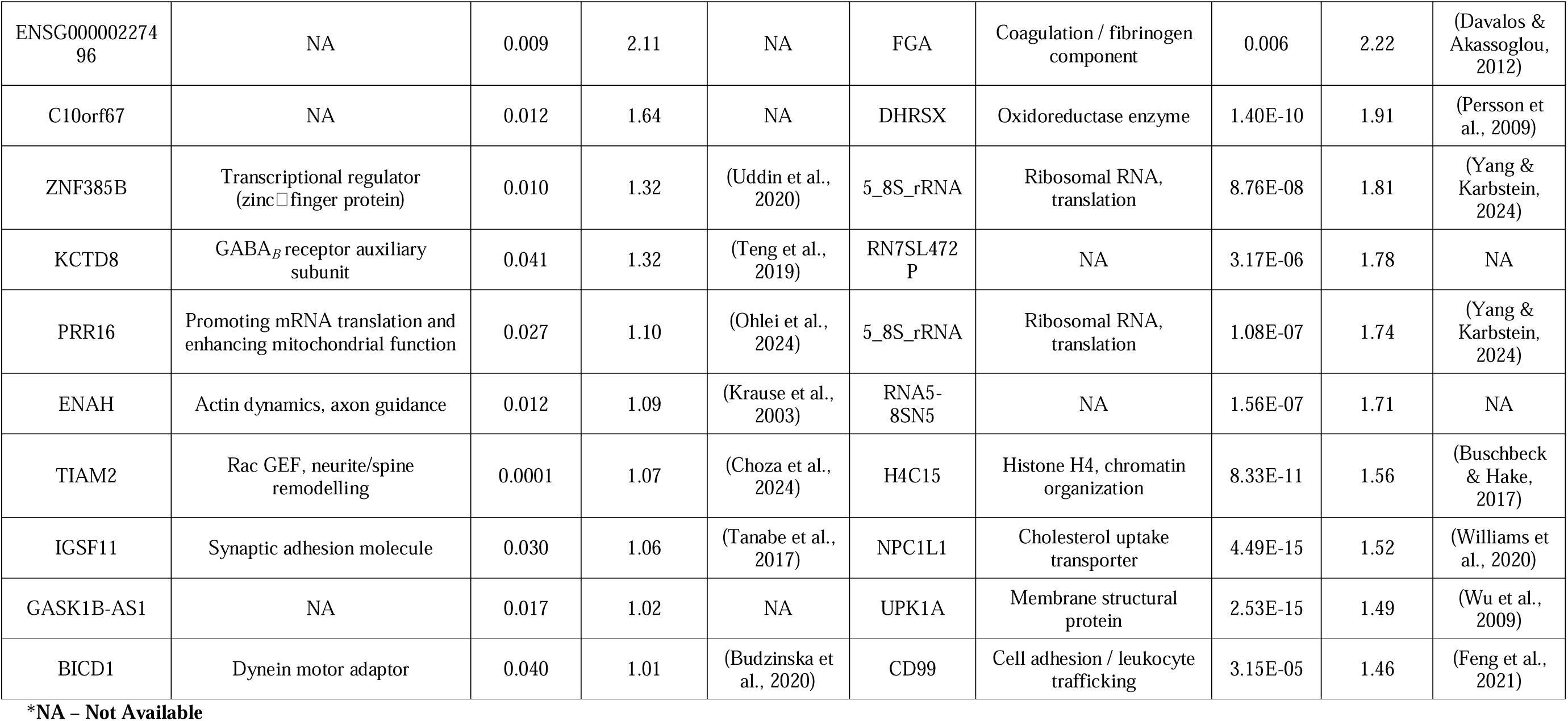
Age-stratified comparison of the most significantly dysregulated genes in the <60 and ≥60 PD cohorts.

*For the* ≥*60 age-group* - DGE testing of 50,489 filtered genes identified 181 significantly dysregulated genes, with 80 downregulated and 101 upregulated (Fig. 4 c)). Top downregulated genes included HOXB13 (-1.998), CHRNA2 (-1.650), SLC35D3 (-1.521), and neuronal markers NPAS4 (-1.442), PVALB (-1.398) (Table 3). Upregulated genes were led by immune transcripts FGA (3.45) and non-coding RNAs (5_8S_rRNA species), reflecting broader immune activation and RNA dysregulation (Kung et al., 2022; Wallings et al., 2020) (Table 3).

#### 3.3.3. Commonly Dysregulated Genes Across Age Groups

To evaluate the consistency of PD-associated transcriptional alterations across age groups and to reconcile differences between pooled and age-stratified analyses, we next examined gene-expression patterns between the <60 and ≥60 cohorts. This step was essential because the complete dataset, driven by a substantially larger ≥60 sample pool (422 ≥60 samples vs 48 <60 samples), may mask early-onset transcriptional signatures, whereas age-restricted subset analysis changes variability and statistical power.

Remarkably, no genes overlapped between the top 20 up/downregulated lists across <60 (31 DEGs) and ≥60 (181 DEGs) cohorts at padj < 0.05 and |log2FC| > 1 (Table 3), confirming age-stratified molecular signatures. The <60 group showed focused set of downregulated novel candidates (LINC02188 -2.296, CALML6 -1.973), while the ≥60 group exhibited broader remodeling characterized by immune activation and neuronal loss.

Cross-validation against the complete dataset further strengthened this pattern: Eight out of ten top downregulated neuronal genes in the complete dataset (PVALB, ARC, NPAS4, FOSB) displayed attenuated or nonsignificant changes in the <60 cohort, indicating that their strong pooled signal was driven largely by the ≥60 cohort (Fig. 4). Conversely, the <60 groups” top candidates (LINC02188, CALML6, TLCD3B) showed minimal changes in the ≥60 group and were therefore, absent from the pooled rankings.

Overall, these results reveal a clear age-dependent divergence, and that PD lacks a single transcriptomic signature spanning all ages. Younger patients show a compact set of novel, stress-related alterations consistent with early calcium/lipid stress, whereas older patients exhibit widespread immune, neuronal, and non-coding RNA dysregulation.

### 3.4. Functional Enrichment Analysis

To explore the biological significance of gene expression changes associated with age in PD, we conducted GO terms and KEGG pathway enrichment analysis. Specifically, we performed enrichment analysis on a) downregulated and upregulated DEGs in the <60 years group and (b) downregulated and upregulated DEGs in the ≥60 years group.

#### 3.4.1. GO analysis for age-based differentially expressed genes

GO enrichment analysis revealed clear age-dependent molecular distinctions between younger (<60 years) and older (≥60 years) individuals with PD.

*For the <60 group -* The analysis showed a predominantly cytoskeleton and neurite-related signature. Upregulated genes were significantly enriched for terms such as filopodium, actin-based cell projection, lamellipodium, dynein complex binding, cytoskeletal anchor activity, and ionotropic glutamate receptor binding (Fig. 5a)). These terms are related to actin-driven structures and cytoskeleton remodeling, thus, suggesting enhanced structural plasticity, motor-protein involvement, and glutamatergic receptor-associated remodeling in younger PD cases, which is consistent with an early compensatory or stress-induced neurite reorganization. On the other hand, a limited set of downregulated GO terms in the <60 group were found, and these mapped to processes such as epidermis morphogenesis, central nervous system neuron axonogenesis, and promoter-specific chromatin binding (Fig. 5a)). This pattern suggests focal suppression of axonogenesis-related and chromatin-regulatory functions, along with an upregulated cytoskeletal remodeling program, supporting a model of early structural instability with focal loss of neuronal growth and transcriptional control.

**Fig. 5.**
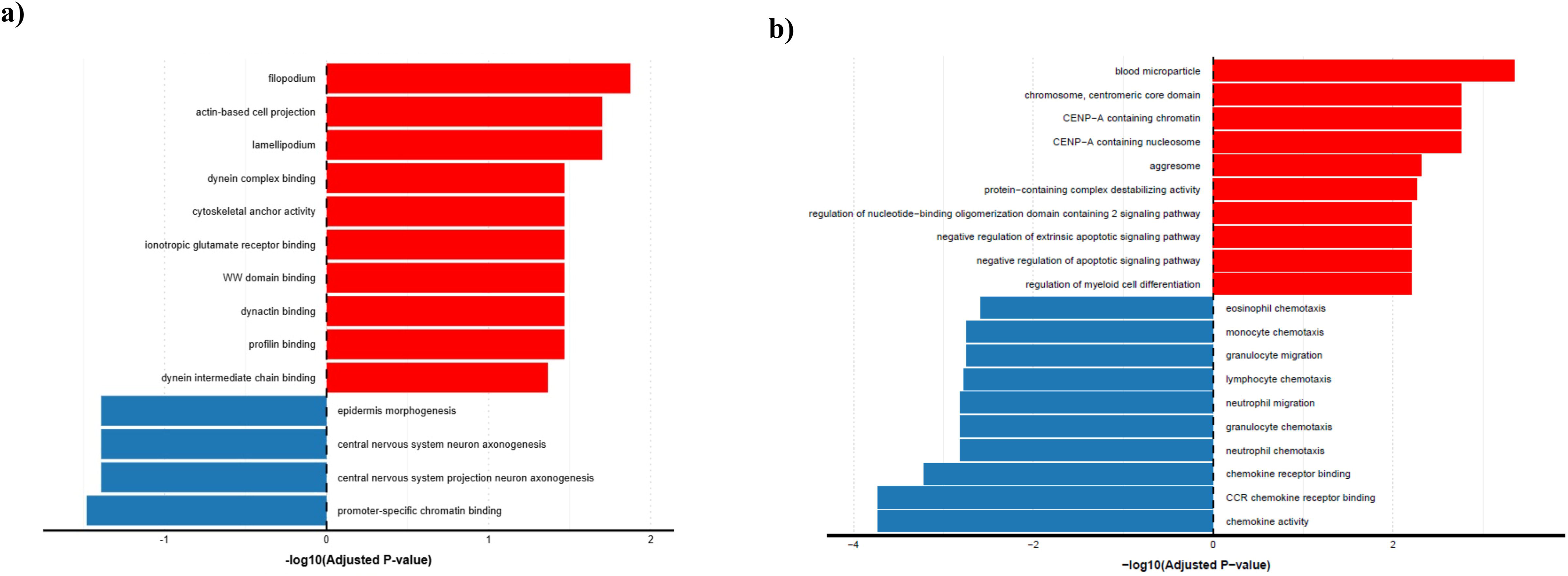
GO enrichment analysis of DEGs in a) individuals aged <60 years, and b) individuals aged ≥60 years. Significantly enriched GO terms are visualized based on their -log10(p-value). Red bars represent GO terms enriched among upregulated genes, while blue bars indicate those enriched among downregulated genes. The top 10 GO terms for each category are displayed, highlighting distinct processes associated with gene expression changes in aging.

*For the* ≥*60 group –* GO analysis showed a distinctly different immune-focused and apoptosis-focused profile as compared to the <60 group. Upregulated genes exhibited a strong enrichment for blood and chromatin-associated processes, including blood microparticle, chromosome, centromeric core domain, CENP-A containing chromatin/nucleosome, aggresome, and protein–containing complex destabilizing activity, as well as pathways regulating apoptotic signaling and myeloid cell differentiation (Fig. 5b)). This suggests elevated circulating/immune-cell activity, centromeric chromatin remodeling, protein quality-control stress, and apoptotic regulation in older PD patients. On the other hand, the downregulated GO terms were focused on chemotaxis and chemokine signaling, including eosinophil chemotaxis, monocyte chemotaxis, granulocyte migration/chemotaxis, lymphocyte chemotaxis, chemokine receptor binding, and CCR chemokine receptor binding (Fig. 5b)). This suggests attenuation of specific leukocyte trafficking and chemokine-receptor interactions, possibly reflecting chronic immune dysregulation in late-onset PD.

Overall, the GO profiles indicate a bias towards cytoskeletal and axonogenic remodelling in younger patients and modulations in blood/immune, chromatin and chemokine signalling in older patients.

#### 3.4.2. KEGG analysis for age-based differentially expressed genes

KEGG pathway enrichment further highlighted distinct age-dependent molecular signatures in PD (Fig. 6).

**Fig. 6.**
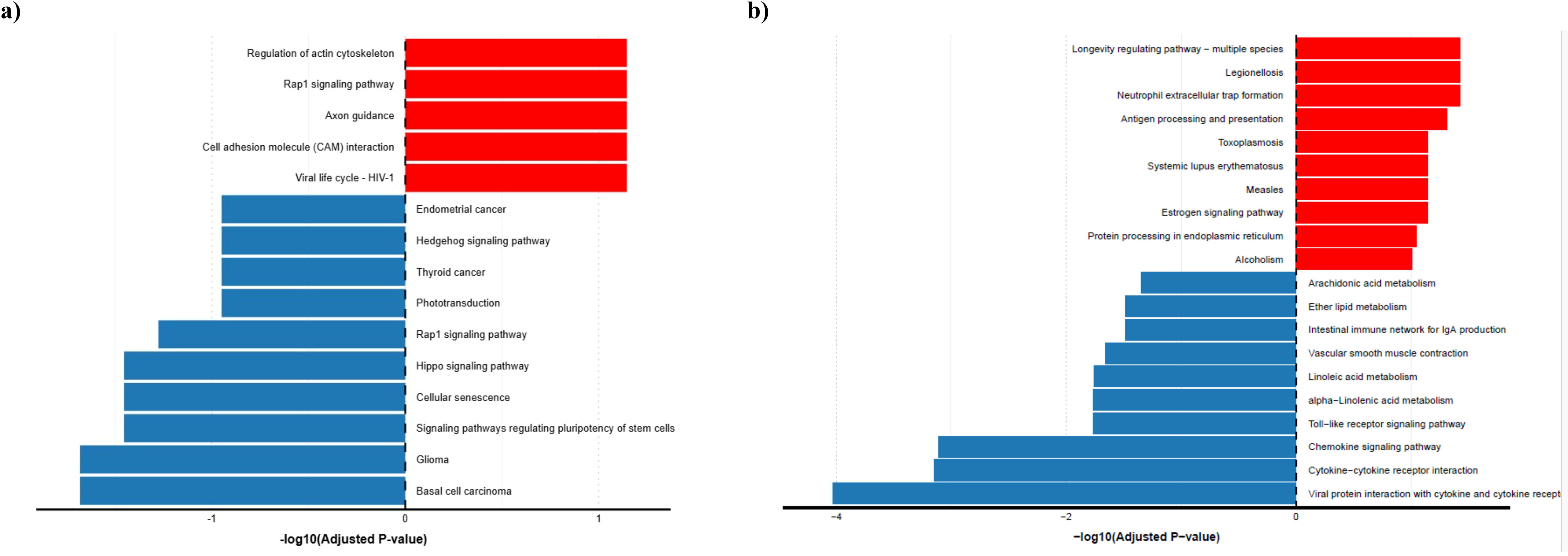
KEGG pathway enrichment analysis of DEGs in a) individuals aged <60 years, and b) individuals aged ≥60 years. Significantly enriched pathways were identified and visualized based on their -log10(p-value). Red bars represent pathways enriched among upregulated genes, while blue bars represent pathways enriched among downregulated genes. The top 10 enriched pathways for each category are shown, highlighting differential pathway activity associated with aging.

*For <60 group -* Upregulated genes were enriched for pathways related to cell adhesion and signaling at the neuronal and immune interface, including regulation of actin cytoskeleton, Rap1 signaling pathway, axon guidance, and cell adhesion molecule (CAM) interaction (Fig. 6a). These pathways collectively implicate dynamic actin remodeling, cell–cell adhesion, and nerve-guidance signaling, consistent with early structural and synaptic reorganization in younger PD cases. Downregulated KEGG pathways in the <60 group were dominated by cancer, developmental, and signaling modules, such as basal cell carcinoma, glioma, signaling pathways regulating pluripotency of stem cells, cellular senescence, Hippo signaling pathway, Rap1 signaling pathway, phototransduction, thyroid cancer, Hedgehog signaling pathway, and endometrial cancer (Fig. 6a). Although many of these are cancer-associated, they converge on core developmental and growth-control signaling (Hippo, Hedgehog, Rap1, pluripotency), indicating selective dampening of proliferative and developmental networks in younger PD, potentially reflecting early-stage restrictions on regenerative or plasticity programs.

*For the* ≥*60 group* - Upregulated genes mapped to pathways such as longevity regulating pathway – multiple species, Legionellosis, neutrophil extracellular trap formation, antigen processing and presentation, toxoplasmosis, systemic lupus erythematosus, measles, estrogen signaling pathway, protein processing in endoplasmic reticulum, and alcoholism (Fig. 6b). These pathways collectively indicate broad activation of innate and adaptive immune mechanisms, pathogen-response modules, ER protein-processing stress, and systemic stress-response signaling in older PD patients. Downregulated KEGG pathways prominently involved lipid and inflammatory mediator metabolism and cytokine signaling, including arachidonic acid metabolism, ether lipid metabolism, intestinal immune network for IgA production, vascular smooth muscle contraction, linoleic and alpha-linolenic acid metabolism, Toll-like receptor signaling pathway, chemokine signaling pathway, cytokine–cytokine receptor interaction, and viral protein interaction with cytokine and cytokine receptor (Fig. 6b). This pattern points to complex remodeling of lipid-derived inflammatory mediators, mucosal immunity, vascular tone, and cytokine/chemokine signaling in late-onset PD, consistent with chronic systemic inflammation and altered immune–vascular cross-talk.

Overall, the KEGG analyses demonstrate that younger PD patients (<60) primarily exhibit alterations in actin cytoskeleton regulation, adhesion, and developmental/growth-control signaling, whereas older patients (≥60) show extensive reconfiguration of immune, pathogen-response, ER stress, and lipid–cytokine signaling pathways.

## 4. Discussion

The age-stratified meta-analysis revealed that PD is associated with distinct, age-dependent transcriptional programs that tightly align with GO and KEGG enrichment patterns across the <60 and ≥60 cohorts. In younger patients, the strongest transcriptomic signals were novel, high-magnitude downregulation of LINC02188, CALML6, MUC20P1, RPS28P7, and TLCD3B, which coincided with GO terms related to axon and neuronal development and with KEGG pathways involving developmental and growth-control signaling. The strongest upregulation was observed for KCTD8, PRR16, TIAM2, and other cytoskeleton-associated genes, which coincided with GO terms linked to dynamic cytoskeletal and protrusive structures, and with KEGG pathways reflecting actin remodeling, neuronal guidance, and adhesion-related signaling. In contrast, the ≥60 group showed strong downregulation of neuronal markers such as NPAS4 and PVALB, which coincided with GO terms capturing impaired leukocyte trafficking and chemokine signaling, and with KEGG pathways tied to disrupted chemokine–cytokine signaling and altered lipid-inflammatory metabolism. The upregulation of immune and non-coding transcripts such as FGA and 5_8S_rRNA species coincided with GO terms related to circulating blood components, chromatin and protein-stress regulation and with KEGG pathways associated with infection and stress responses. Together, the analyses indicate that younger patients show a focused signature of cytoskeletal and axonogenic remodeling, while older patients display broad immune, chromatin, and metabolic dysregulation, establishing age as the primary axis of molecular heterogeneity in PD.

To put these age-dependent signals into context, we compared the stratified results with the pooled meta-analysis. Canonical neuronal activity–dependent genes that dominate prior PD transcriptomic reports, such as PVALB (Lanoue et al., 2013; Zhang, B. et al., 2024), ARC (Calabresi et al., 2009; Garcia et al., 2017), NPAS4 (Mousa et al., 2023), and FOSB (Ebihara et al., 2011), are strongly downregulated in the complete dataset and in the ≥60 cohort, consistent with the longstanding “global neuronal suppression” signature described across substantia nigra, cortex, and blood (Grünblatt et al., 2004; Irmady et al., 2023; Moran et al., 2006). However, these genes lose statistical significance or show attenuated effect sizes in the <60 group, revealing that the established PD neuronal-suppression phenotype is largely an age-amplified, ≥60-driven signal rather than a universal PD hallmark. Conversely, early-onset-enriched candidates such as LINC02188, CALML6, and TLCD3B show large fold changes exclusively in the <60 group, suggesting age-specific or early-stage biological processes that are diluted or obscured in pooled analyses. This cross-age behavior implies that suppression of activity-regulated neuronal gene programs reflect late-stage, age-amplified PD, while younger-onset PD is characterized by lncRNA-, calcium-, and lipid-linked structural remodeling that may precede interneuron loss.

These findings differ from previous PD meta-analyses, which consistently report synaptic dysfunction, mitochondrial impairment, and immune activation as hallmark molecular PD signatures, but have seldom targeted age as an independent factor. Earlier bulk RNA-seq and microarray meta-analyses of substantia nigra and cortical tissue have reported age-invariant downregulation of synaptic pathways accompanied by increased inflammatory activation (Moran et al., 2006; Simunovic et al., 2009; Sutherland et al., 2009). Recent meta-analyses integrating brain and blood transcriptomic data along with network-based approaches, have highlighted immune, oxidative phosphorylation, and proteostasis pathways as common PD signatures (Kong et al., 2018; Kurvits et al., 2021; Navarrete et al., 2025). However, these studies didn”t segregate the patients on the basis of age, thus, providing limited information on molecular features associated with early- and late-onset disease (Kong et al., 2018; Kurvits et al., 2021; Navarrete et al., 2025). Similarly, several meta-analyses studies exploring PD-related genes and pathways have identified age-invariant modules related to synaptic transmission, mitochondrial dysfunction, and neuroinflammation (Chen et al., 2022; Wang et al., 2017; Zhou et al., 2023). This limitation was addressed in the current study by directly stratifying cohorts by age and integrating GO/KEGG enrichment with cross-validation against the pooled dataset. The analysis showed that the widely recognized PD transcriptional signature is largely driven by older patients. Early-onset PD displayed a much more focused molecular profile dominated by cytoskeletal organization and axonal processes, indicating biological changes that are largely masked when age groups are analyzed together. Overall, these findings suggest that aging substantially influences the transcriptional landscape of PD and that the molecular mechanisms underlying the disease vary with age (Collier et al., 2017; Hou et al., 2019; Reeve et al., 2014).

A key question arising from these findings is whether the observed dysregulation is driven by normal aging, or PD pathology, or their combination. Previous studies have shown that healthy aging is generally accompanied by alterations in immune and mitochondrial pathways, while neuronal activity-dependent genes, such as ARC and NPAS4, progressively decline across the cortex and hippocampus, contributing to age-related impairments in synaptic function and cognition (Hou et al., 2019; Misrani et al., 2021). Accordingly, the significant downregulation of PVALB, NPAS4, and ARC observed in the ≥60 cohort is likely to reflect the combined effects of age-associated alterations and PD-related neurodegeneration (Hou et al., 2019; Shepard et al., 2019). In contrast, several significantly altered genes in the <60 group, including LINC02188, and TLCD3B, have not been linked to normal brain aging. Also, calcium□regulating transcripts such as CALML6 have been associated with PD risk and dopamine□neuron sensitivity and not with normal aging (Surmeier et al., 2010; Surmeier et al., 2011). These observations suggest that the molecular changes detected in early-onset PD primarily represent disease-specific mechanisms occurring within a relatively intact neuronal environment.

Despite its strengths, this study has several limitations. First, the <60 cohort is relatively small (36 controls, 12 PD), reducing power to detect subtle changes and likely underestimating the breadth of early-onset dysregulation. Second, bulk RNA-seq limits cell-type resolution, making it difficult to disentangle interneuron, microglial, and peripheral immune contributions. Third, integrative analyses across heterogeneous tissues and sequencing platforms introduce residual confounders such as medication exposure, disease duration, and comorbidities. Finally, functional inference from GO/KEGG relies on existing annotations and may underrepresent emerging lncRNAs and pseudogenes such as LINC02188 and RPS28P7; experimental follow-up is required to establish their functional or biomarker relevance.

These limitations underscore the need for age-aware experimental and clinical frameworks. Future work should integrate age with tissue and region specificity across blood, CSF, gut, skin, and distinct brain regions to identify circulating signatures that mirror or precede central pathology. Longitudinal, multi-omic datasets-including miRNA, lncRNA, lipidomic, and single-cell profiles-will be essential to determine whether the age-stratified signatures identified here predict progression from prodromal to overt PD and to separate region-specific from systemic disease processes.

## 5. Conclusion

This age-stratified meta-analysis shows that Parkinson”s disease lacks a unified transcriptomic signature across age groups. Younger patients (<60 age) display a focused pattern of cytoskeletal and axonogenic remodeling with suppression of developmental pathways, suggesting early structural vulnerability. Older patients (≥60 age) show broad immune activation, chromatin and protein-stress remodeling, disrupted lipid–cytokine signaling, and dominant neuronal loss signatures. The strong, age-specific signals identified here highlight potential early-onset biomarkers such as LINC02188, CALML6, TLCD3B, MUC20P1 and RPS28P7, and late-onset candidates such as NPC1L1, and IL3RA. We propose these potential biomarkers for further experimental and longitudinal validation.

## Supporting information

Supplementary Sheet S1

Supplementary Sheet S2

Supplementary Sheet S3

Supplementary Sheet S4

Supplementary Sheet S5

Supplementary Sheet S6

## Acknowledgements

The authors would like to thank the scientific community.

## Funding

This research did not receive any specific grant from funding agencies in the public, commercial, or not-for-profit sectors.

## Competing interests

The authors have no relevant financial or non-financial interests to disclose.

## Authors Contributions

Data Curation, Formal analysis, Writing - original draft preparation, and investigation: **Prakashini Saroj Nilgirwar**; Writing - original draft preparation, Writing - review and editing, and investigation: **Sahil Jain**; Data Curation and Formal analysis: **Bhakti Badhe**; Data curation and Formal analysis: **Rituja Shinde**; Writing - review and editing: **Manoj Baranwal**; Conceptualization, Methodology, Supervision: **Dimple Davray**

## Notes

### Competing Interest Statement

The authors have declared no competing interest.

